# Delineating the Genetic Component of Gene Expression in Major Depression

**DOI:** 10.1101/2020.03.24.004903

**Authors:** Lorenza Dall’Aglio, Cathryn M. Lewis, Oliver Pain

## Abstract

**Background:** Major Depression (MD) is determined by a multitude of factors including genetic risk variants which regulate gene expression (GE). Here, we examined the genetic component of GE in MD by performing a Transcriptome-Wide Association Study (TWAS), inferring GE-trait relationships from genetic, transcriptomic and phenotypic information.

**Method:** Genes differentially expressed in depression were identified with the TWAS FUSION method, based on summary statistics from the largest genome-wide association analysis of MD (*N_cases_* = 135,458) and GE levels from 20 tissue datasets. Follow-up analyses were performed to extensively characterize the identified associations: colocalization, conditional, and fine-mapping analyses together with functionally-enriched pathway investigations.

**Results:** Transcriptome-wide significant GE differences between cases and controls were found at 91 genes, 50 of which were not found in previous MD TWASs. Of the 91 significant genes, eight represented strong, colocalized, and potentially causal associations with depression, which were independent from the effect of nearby genes. Such “high-confidence associations” include *NEGR1, CTC-467M3.3, TMEM106B, CTD-2298J14.2, CCDC175, ESR2, PROX2, ZC3H7B.* Lastly, TWAS-based enrichment analysis highlighted dysregulation of gene sets for long term potentiation, dendritic shaft, and memory processes in MD.

**Conclusion:** This study has shed light on the genetic component of GE in depression by characterizing the identified associations, unravelling novel risk genes, and determining which associations are congruent with a causal model. These findings can be used as a resource for prioritizing and designing subsequent functional studies of MD.

## Introduction

Major depression (MD) constitutes one of the largest contributors to global disability worldwide, affecting around 322 million people and accounting for approximately 50 million years lived with disability (1). This disorder is of complex origin, being determined by the interplay of a multitude of environmental factors (e.g. life events) and genetic variations (2). The genetic contribution to MD has been confirmed by twin studies, which yielded heritability estimates of approximately 37% (3), and genome-wide association studies, which estimated heritability of 9% captured by common single nucleotide polymorphisms (SNPs) (4).

Genome-wide association studies (GWASs) have started to identify the specific genomic loci underlying depression genetic risk. In one of the most recent and largest GWASs to date (*N_cases_ = 135,458*), Wray et al. (5) uncovered 44 independent loci associated with MD. Drawing on a larger sample (*N_cases_ = 246,363*) and a broader phenotype definition, Howard et al. (4) observed 102 independent loci, further demonstrating the polygenicity of the disorder.

While genome-wide studies have enabled the identification of SNPs conferring susceptibility to depression, the *functional significance* of such genetic variants remains to be elucidated. The examination of intermediary processes between genomics and the phenotype, such as gene expression (GE), would permit greater insight into the molecular mechanisms underlying depression.

GE is a biological process permitting the translation of the genetic code into functional products, such as proteins and functional RNA (e.g. transfer RNA). While GE is affected by extrinsic exposures such as diet (6), a substantial role for SNPs on the transcriptome has also been observed, with 50% of common variants being associated with GE, in any tissue (7). Genetic variants affecting GE are referred to as Expression Quantitative Trait Loci (eQTL).

Studies mapping eQTLs enabled the development of publicly available reference panels containing information on SNP – GE relationships, in multiple tissues across different samples. Based on these, the *cis*-genetic component of GE can be predicted in any genetically mapped trait. This allows for the analysis of GE – trait associations with larger samples, inexpensively, and in multiple tissues, as opposed to classical observed GE studies. Importantly, since such novel methods solely capture the genetic component of GE, differential expression that is a consequence of depression (reverse causation) can be excluded.

Several methods to predict GE – trait associations are available. These leverage three key sources of information: *(i)* reference panels with data on SNPs – GE associations (e.g. the Genotype-Tissue Consortium (GTEx)), *(ii)* individual- or summary statistic-level GWAS data capturing SNP-trait associations, and *(iii)* a linkage disequilibrium (LD) reference. Of the available methods, Summary-data based Mendelian Randomization (SMR) (8) and Transcriptome-Wide Association Studies (TWASs) (9,10) are the most commonly used. While the GE – trait association is explored SNP-by-SNP in SMR, TWASs entail a gene-by-gene approach. The integration of SNP effects into genes lowers the multiple testing burden and increases power, thus allowing TWASs to identify genetic associations potentially overlooked by SMR studies and GWASs (9).

To date, four studies have tested for an association between the genetic component of GE and depression (5,11–13). Wray et al. (5) showed transcriptomic dysregulation in MD at 17 genes in the dorsolateral prefrontal cortex (DLPFC). Other studies extended these findings by identifying a greater number of associations, within major (12,13) as well as broad depression (11), across multiple tissues (i.e. distinct brain tissues and whole blood). In these studies, three genes (*NEGR1, LRFN5,* and *ESR2*) were highlighted due to their involvement in the production of proteins targeted by psychotropic drugs, indicating that such loci may constitute potential therapeutic targets (12).

These studies have shed further light on the genetic and functional mechanisms of depression, and in this study we extensively examine the characteristics of the identified TWAS associations by exploring *(i)* the origin of TWAS gene associations (pleiotropic SNP effects on GE and depression vs. linkage), *(ii)* co-expression patterns among significant genes, *(iii)* potentially causal effects through fine-mapping procedures, as well as *(iv)* functionally-enriched pathways in depression. These analyses enable key genes for depression to be highlighted, providing novel insights into the biology of the disorder. This study extends the tissues used for transcriptome-wide studies of depression, beyond brain and blood transcriptomic profiles to the Hypothalamus-Pituitary-Adrenal gland (HPA) / Thyroid (HPT) axes. These tissues are important in MD due to their involvement in the hypo/hyperactivity of stress responses, sleeping difficulties, and weight loss, which are physiological dysfunctions shown in depressed patients (14–16).

Overall, this study aims to provide a comprehensive and in-depth characterization of the association between the *cis-*genetic component of gene expression and major depression through comprehensive analyses within a multitude of tissues.

## Methods and Materials

### Datasets

The analyses used *(i) genome-wide* summary statistics from the Wray et al. (5) GWAS of MD, *(ii)* 20 SNP-weight sets from four separate transcriptomic reference samples, and *(iii)* the 1000 Genomes Project reference for LD estimation.

Firstly, we leveraged summary statistics from the Psychiatric Genomics Consortium MD GWAS (Wray et al. (2018)), which includes information on the genetic susceptibility to MD from seven samples: the PGC studies, deCODE, Generation Scotland, GERA, iPSYCH, the UK Biobank, and 23andMe. The total sample included 135,458 MD cases and 344,901 controls and summary statistics provided association results from 1,185,038 HapMap3 SNPs.

Secondly, SNP-weights from distinct tissues and samples were used. *SNP-weights* represent the correlations of SNPs with the expression of their annotated gene (10). SNP-weights from *postmortem* brain tissue, whole blood, peripheral blood, and the adrenal, pituitary and thyroid glands were downloaded from the TWAS FUSION website (see URLs). The weights pertained to four RNA reference samples: *(i)* Netherlands Twins Register (NTR) and *(ii)* Young Finns Study (YFS), which both provide information on blood tissue gene expression, *(iii)* CommonMind Consortium (CMC), which assessed the DLPFC with two distinct assays, and *(iv)* the Genotype-Tissue Expression (GTEx) Consortia, a state-of-the-art study where expression in multiple brain and peripheral tissues (e.g. blood, thyroid) was measured (10,17,18). Of the GTEx data, we used SNP-weights for 16 tissues from brain, thyroid, pituitary and adrenal glands, and blood. We use the term *SNP-weight sets* for SNP-weights within a given sample and tissue (e.g. GTEx thyroid, NTR peripheral blood). Each gene within a given SNP-weight set constitutes a *feature* or *gene-tissue pair.* SNP-weight set characteristics, including the RNA reference sample (e.g. GTEx) and the assay used (e.g. RNA-seq), are available in Supplementary Table S1. The 1000 Genomes Phase 3 European LD reference (*N*=489) (19) was downloaded from the FUSION website (see URLs).

### Statistical Analyses

All statistical analyses were performed in Bash or R (version 3.5.0).

#### Transcriptome-wide significance threshold

To compute the transcriptome-wide significance threshold for this study, we leveraged a permutation procedure used in a previous TWAS study (20) (Supplementary Text). This approach estimates a significance threshold adjusted for the number of features tested in the study, taking into account the correlation between features within and across SNP-weight sets. The threshold for transcriptome-wide significance was *p*=2.13×10^-6^ (for a false positive rate of *α* = 0.05), with a more stringent significance threshold of *p*=4.38×10^-8^, for α=0.001.

#### TWAS FUSION and colocalization

A TWAS FUSION analysis was run on autosomal chromosomes, following the TWAS FUSION protocol with default settings (Supplementary Text) (see URLs). Colocalization was assessed using the *coloc* R package for all genes meeting transcriptome-wide significance, and within a 1.5 Mb window. This Bayesian approach estimates the posterior probability that associations within a locus for two outcomes are driven by a shared causal variant. It thus enables the distinction between associations driven by horizontal pleiotropy (one causal SNP affecting both GE and MD; posterior probability PP4) and linkage (two causal SNPs in LD affecting GE and MD separately; posterior probability PP3). More details are available in the Supplementary Text.

#### Conditional analysis

Genomic regions with significantly associated features were further analysed with a conditional analysis to determine whether multiple associated features within a given locus represent multiple independent associations, or a single association due to correlated predicted expression between features. This was performed using the FUSION software, which jointly estimates the effect of all significant features within each locus by using residual SNP associations with depression after accounting for predicted expression of other features. This process identifies which features represent independent associations (termed jointly significant), and which features are not significant when accounting for the predicted expression of other features in the region (termed marginally significant) (10) (Supplementary Text). We additionally calculated to what extent the GWAS associations within each locus could be explained by functional associations detected in this TWAS (i.e. the variance explained) (Supplementary Text).

#### TWAS Fine-mapping

FOCUS, a TWAS association fine-mapping method, was utilized to identify which features are likely to be causal within regions of association (21). Analogous to statistical fine-mapping of GWAS results, FOCUS estimates the posterior inclusion probability (PIP) of each feature being causal within a region of association, using the sum of posteriors to define the default 90% credible set, a set of features likely to contain the causal feature. The method includes a null model, where the causal feature is not present. The PIP of individual features is also of interest, with a PIP>0.5 indicating a feature is more likely to be causal than any other feature in the region. The FOCUS finemap function was then applied *without* the tissue prioritisation option.

#### TWAS-based gene set enrichment analysis

Finally, we used a previously applied approach for TWAS-based gene set enrichment analysis (TWAS-GSEA) (20) to identify functionally-informed dysregulated pathways characterizing MD (Supplementary Text). A linear mixed model was used to test for an association between *z*-scores indicating non-zero association for each feature and gene set membership, while adjusting for gene length and numbers of SNPs within the gene region, and accounting for correlation between features. Linear mixed models were fitted using the R package *lme4qtl* (22). TWAS-GSEA used 7,246 hypothesis-free gene sets from MSigDB (v6.1) and 76 candidate gene sets from (5), which included FMRP binding partners, de novo mutations, GWAS top SNPs, and ion channels. A minimum of five genes within the gene set was required to perform the TWAS-GSEA analysis. TWAS-GSEA was run using different TWAS results to identify gene-sets enriched across all tissues, within tissue groups (brain, blood, HPA, HPT), and within each SNP-weight set. If multiple features for a single gene were present, the feature explaining the largest amount of variance in the gene’s expression was retained. Multiple testing correction for the number of gene-sets tested was performed using the FDR method separately for candidate and hypothesis-free analyses, using a *q*-value of 0.05 to determine significance.

#### Comparisons with previous literature of observed and predicted gene expression

Lastly, we compared our findings with previous literature of *observed* and *predicted* gene expression. The former comparison was performed to evaluate to what extent we could replicate the findings from the largest transcriptomic study of depression to date (*N* = 1,848) (6). This assessed whole blood gene expression using microarray technology in the Netherlands Study of Depression and Anxiety. Genes were considered as replicated if Bonferroni significant (*p* < .05/number of unique genes of nominal significance) in our study. Consistent direction of effect was not a requirement for replication.

We additionally contrasted our results to three previous studies (5,12,13) of predicted GE in MD to evaluate the consistencies across studies and our novel contributions. Overall, we considered 133 unique genes, significant in *any* SNP-weight set, in *either* of the four TWASs. Genes identified as statistically significant by this, but not previous TWASs, were labelled as novel.

## Results

### TWAS Analysis

We identified 177 significant features, from 91 unique genes, which were differentially expressed *(p* < 2.13×10^-6^) across multiple SNP-weight sets (Figure 1, Supplementary Figure 1a- b; Supplementary Table S2). Of these gene-tissue pairs, 95 were upregulated while 82 were downregulated. The most significant feature was *NEGR1* (GTEx whole blood) (*z* = 8.76, *p* = 1.94×10^-18^). In total, 41 unique genes (from 76 features) were located more than 500 kb from a genome-wide significant SNP in the MD GWAS, a criteria previously used to indicate a novel association (23). The largest number of associations were from the GTEx Thyroid SNP-weight set (23 associated genes), but inferences on tissue enrichment are difficult as SNP-weight sets differ in their characteristics such as sample size, and the number of tested features. Whilst 36 of the 91 unique genes showed transcriptomic alterations across multiple SNP-weight sets, 55 genes were up-/down-regulated in only one SNP-weight set. Of note, 64 associations (from 29 unique genes) were within the extended Major Histocompatibility Complex (MHC) region (chromosome 6: 26-34 Mb). This region is gene rich and characterized by extensive LD, so these associations should be interpreted with caution.

**Figure 1.**
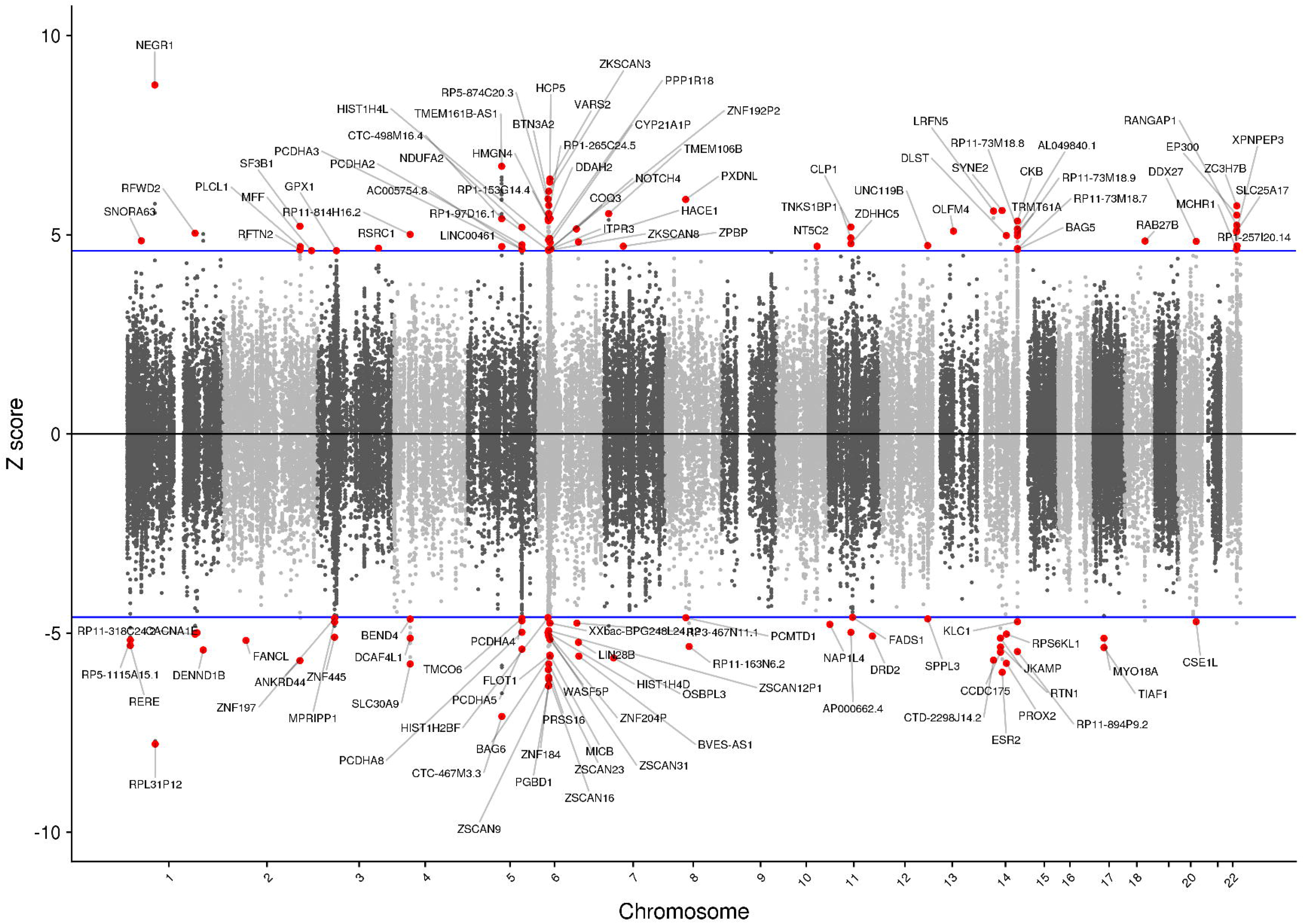
The relationship between gene expression and Major Depression: Manhattan-style plot of *z*-scores for each of the tested genes, across all autosomes and tested SNP-weight sets. Blue lines indicate the transcriptome-wide significance threshold. The names of statistically significant genes are shown.

### Colocalization

We evaluated the colocalization status of a gene by calculating the posterior probability that the genetic and functional associations derived from (i) *distinct* causal SNPs (PP3), and *(ii)* a *shared* causal SNP (PP4) (24). Of the 177 significant features, 101 (57 unique genes) were considered as colocalized based on their high PP4 (> 0.8), in line with previous literature (25,26) (Supplementary Table S3). The strongest posterior probabilities for colocalization were for *FLOT1, RP5-1115A15.1,* and *CKB* (PP3 = 0, PP4 = 1).

### Conditional Analysis

We observed that multiple significant features resided within the same locus (defined as a 1.5 Mb window), for a total of 35 genomic regions. Conditional analysis of the 177 transcriptome-wide significant features identified 47 jointly and 130 marginally significant features, from 44 and 47 unique genes respectively.

We observed anomalous results in the MHC region (Supplementary Table S4) likely reflecting technical problems due to the mismatch between the LD references used in the different samples (GWAS and functional data) and were thus excluded. Conditional analysis results for each locus are shown in Table 1, with marginally significant features shown in Supplementary Table S5.

**Table 1.**
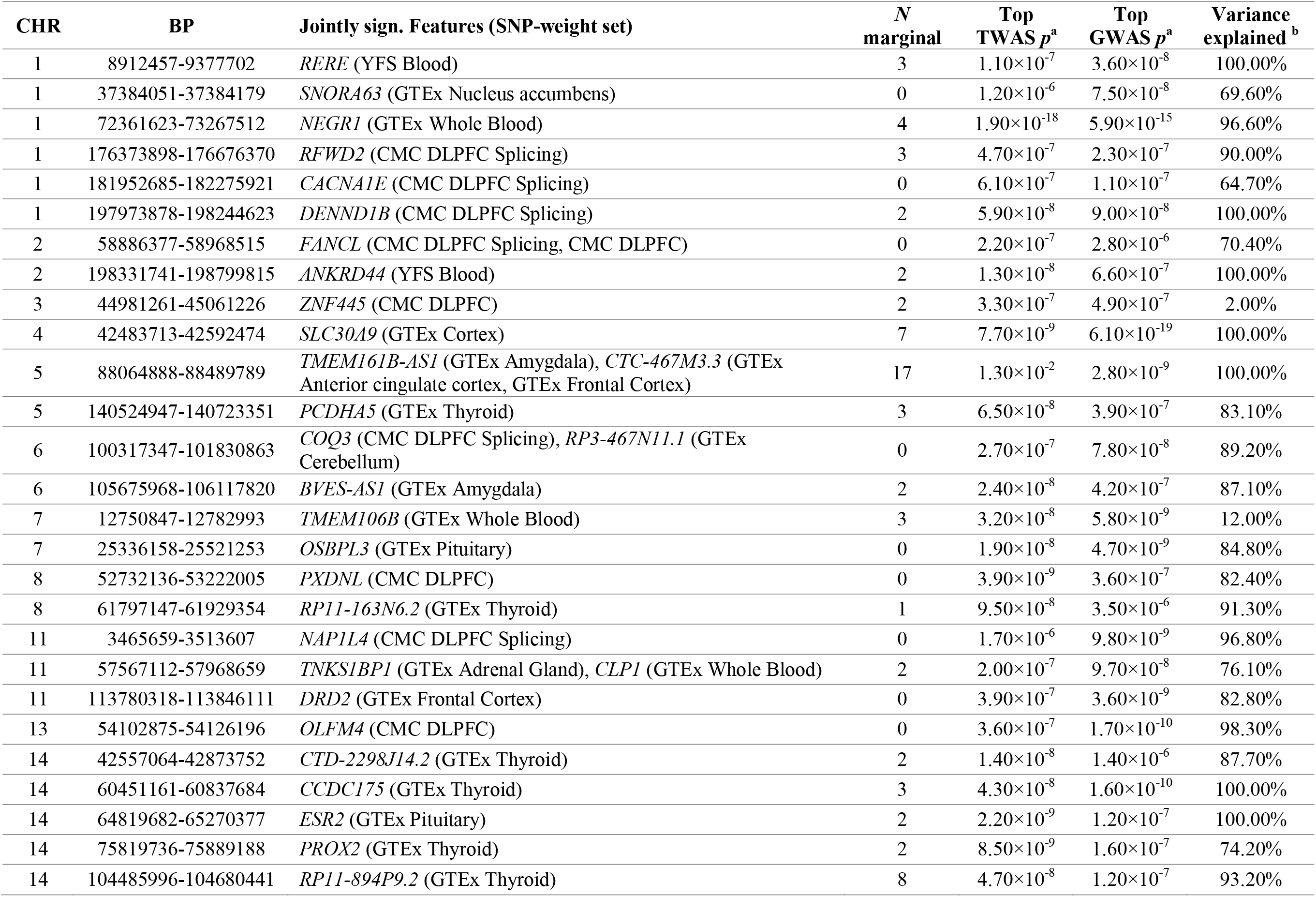
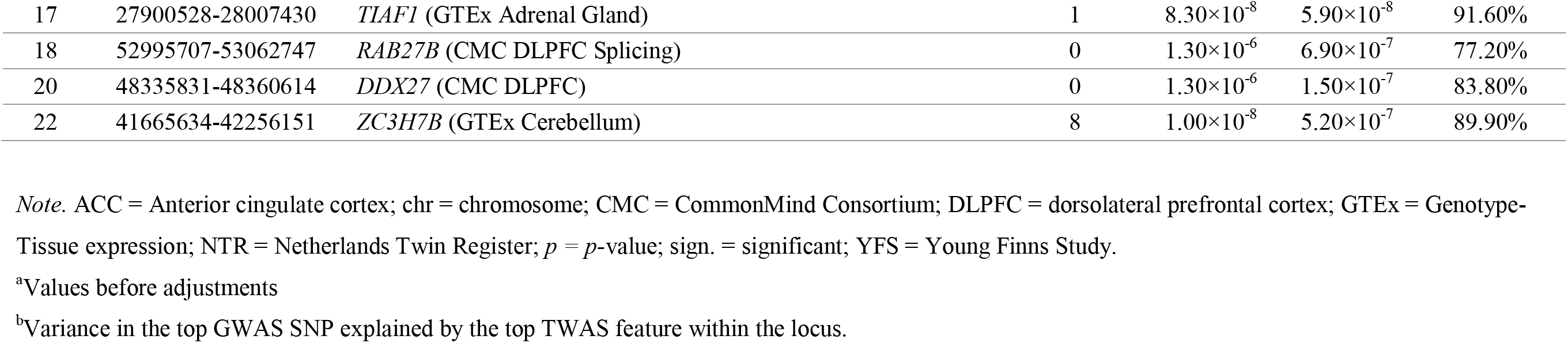
Conditional Analysis Findings: Jointly and Marginally Significant Features per Locus

We additionally investigated the effect of adjusting for features’ gene expression on associations between SNPs and the trait (i.e. GWAS findings). The median GWAS *p-*value before conditioning (*p*=9.7×10^-8^) reduced to *p*=0.078 following adjustments, showing that GWAS associations were mostly accounted for by TWAS significant features. The variance explained was highly variable (median variance explained = 89%, range = 2% – 100%). Of note, genome-wide associations in seven loci were fully explained by our TWAS results (Table 1). This included a locus with *ESR2* (GTEx Pituitary) as the jointly significant feature accounting for GWAS signal. Instances where the TWAS association does not explain the most significant GWAS association in the locus can occur when the TWAS associations are driven by a secondary independent GWAS association.

### Statistical fine-mapping

FOCUS was performed for all loci containing transcriptome-wide significant associations to calculate the probability estimates of causality (PIP) for each feature, and define 90% credible sets (Supplementary Tables 6). Of the 17 loci containing multiple transcriptome-wide significant genes, FOCUS identified 6 loci containing only one significant gene in the 90% credible set, demonstrating FOCUS’s ability to pinpoint a specific causal gene. Two clear examples of this are GTEx Whole blood *NEGR1* expression (PIP = 1) and GTEx Thryoid *PROX2* expression (PIP = 0.98), indicating they are likely causal for the associations with depression. The significance of TWAS associations is not the only consideration when fine-mapping likely causal features, and as a result within two of the 33 genomic loci containing significant TWAS associations excluding the MHC region, the feature with the highest PIP was not transcriptome-wide significant *(ASTN1* for chr1:176373898-176676370 (TWAS.Z = −3.96, PIP = 0.705), and *POLI* for chr18:51554436-18:55213381 (TWAS.Z = −4.3408, PIP = 0.225)).

### “High-Confidence” Associations

In a data-driven approach, we highlighted the genes that are most relevant to MD. Firstly, in the TWAS results, we applied a more stringent significance threshold of *p* < 4.38×10^-8^ (*α* = 0.001) to minimize the chance of false positives. Secondly, from these highly significant genes, we identified those which were colocalized (PP4 > 0.8) and likely to be causal (PIP estimate of >0.5). This strategy showed eight “high-confidence” associations (Table 2). The *NEGR1* gene constituted the most significant hit with the highest posterior probability of colocalization and of causality (PP4=0.993; FOCUS PIP=1.00). All features highlighted using this approach were jointly significant in the conditional analysis, meaning they are independently associated with depression, and are the most significant features within each region of association.

**Table 2.** Characteristics of the High-Confidence Associations: Highly Significant, Colocalized, and Independent Features

| Location | Gene | SNP-weight set | Coloc PP3<sup>a</sup> | Coloc PP4<sup>b</sup> | Z-score<sup>c</sup> | P-value<sup>c</sup> | PIP<sup>d</sup> |
| --- | --- | --- | --- | --- | --- | --- | --- |
| chr1:71861623-72748417 | *NEGR1* | GTE<sub>Ex</sub> Whole Blood | 0.007 | 0.993 | 8.761 | 1.94×10<sup>-18</sup> | 1 |
| chr5:87988462-87989789 | *CTC-467M3.3* | GTE<sub>Ex</sub> Frontal Cortex | 0.035 | 0.85 | -7.092 | 1.33×10<sup>-12</sup> | 0.867 |
| chr7:12250867-12282993 | *TMEM106B* | GTE<sub>Ex</sub> Whole Blood | 0.008 | 0.991 | 5.531 | 3.18×10<sup>-8</sup> | 0.794 |
| chr14:42057064-42074059 | *CTD-2298J14.2* | GTE<sub>Ex</sub> Thyroid | 0.022 | 0.978 | -5.679 | 1.36×10<sup>-8</sup> | 0.563 |
| chr14:59971779-60043549 | *CCDC175* | GTE<sub>Ex</sub> Thyroid | 0.018 | 0.979 | -5.479 | 4.28×10<sup>-8</sup> | 0.543 |
| chr14:64550950-64770377 | *ESR2* | GTE<sub>Ex</sub> Pituitary | 0.026 | 0.86 | -5.982 | 2.20×10<sup>-9</sup> | 0.592 |
| chr14:75319736-75330537 | *PROX2* | GTE<sub>Ex</sub> Thyroid | 0.02 | 0.962 | -5.758 | 8.51×10<sup>-9</sup> | 0.984 |
| chr22:41697526-41756151 | *ZC3H7B* | GTE<sub>Ex</sub> Cerebellum | 0.031 | 0.862 | 5.729 | 1.01×10<sup>-8</sup> | 0.532 |

*Note.* Chr = chromosome; Coloc = Colocalization; CMC = CommonMind Consortium; GTE<sub>Ex</sub> = Genotype-tissue expression; NTR = Netherlands Twin Register; YFS = Young Finns Study.
<sup>a</sup>PP3 = Probability of a GWAS and TWAS association, at distinct causal SNPs.
<sup>b</sup>PP4 = probability of a GWAS and TWAS association, at shared causal SNPs.
<sup>c</sup>TWAS association results.
<sup>d</sup>Posterior Inclusion Probability estimated by FOCUS.

### TWAS-GSEA

TWAS-based gene set enrichment analysis identified 15 gene sets in the candidate analysis, and nine hypothesis-free gene-sets, when either combining TWAS results across brain SNP-weight sets, or when using individual SNP-weight sets (Tables 3 and 4). Of note, the hypothesis-free analysis showed genes differentially expressed in the brain of depressed individuals were significantly enriched for functions relating to long-term potentiation of synapses, the dendritic shaft of neurons, and memory. No significant associations were detected when combining results across all SNP-weight sets or when combining SNP-weight sets for blood, HPA, or HPT tissue groups.

**Table 3.**
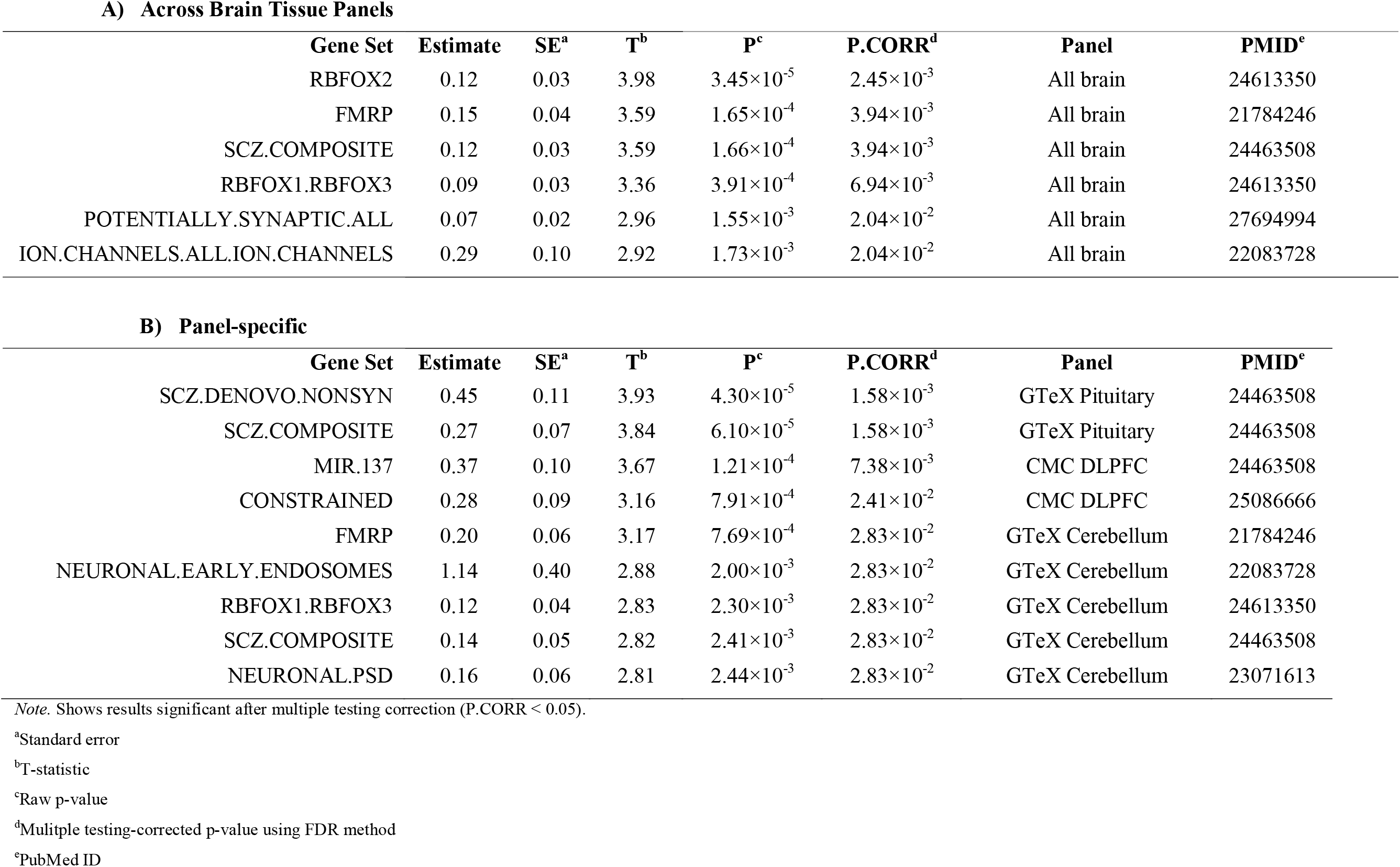
Candidate Gene Set Enrichment Results based on TWAS results from A) all brain tissue panels, and B) each panel separately.

**Table 4.**
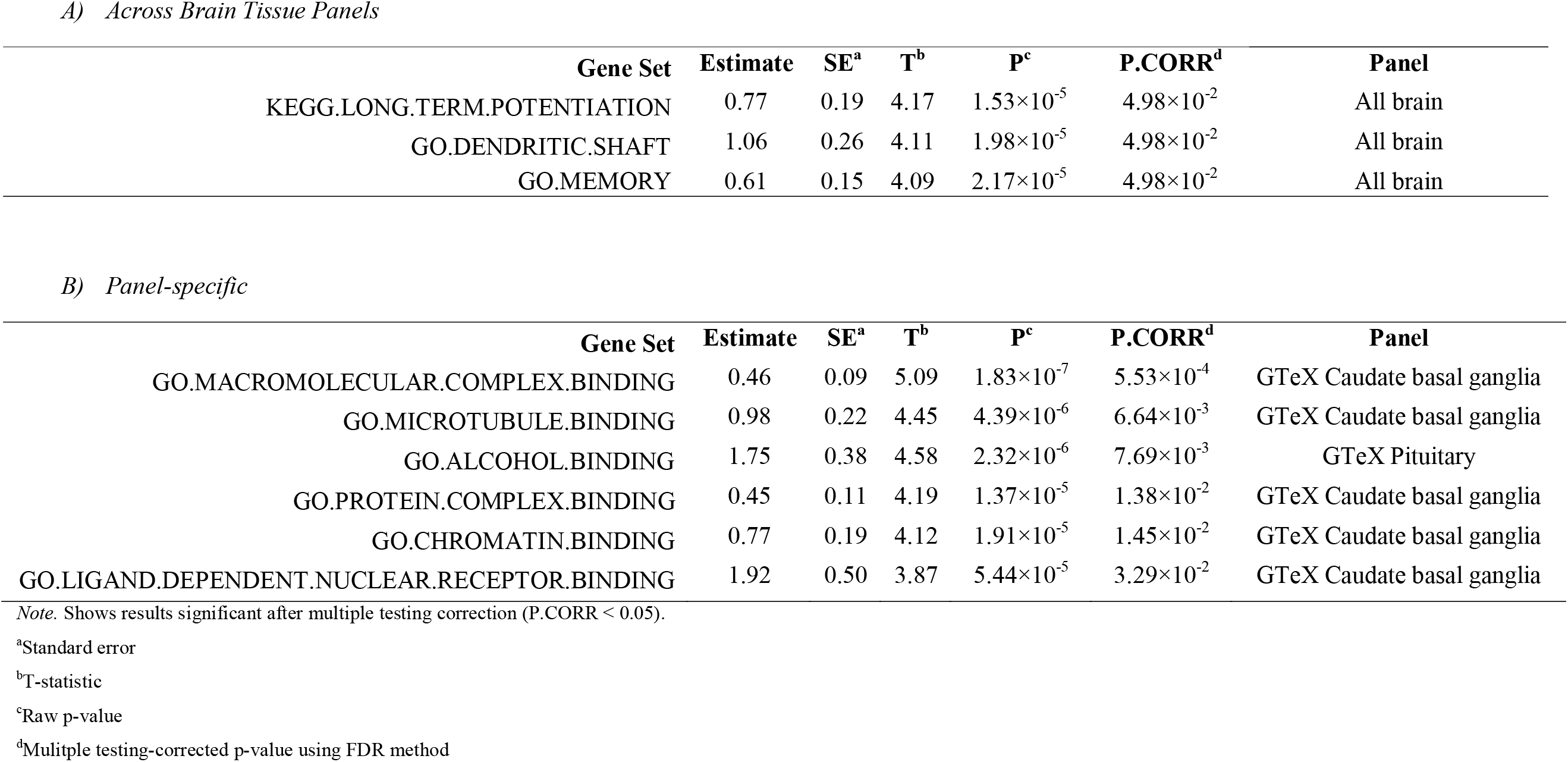
Hypothesis-free Gene Set Enrichment Results based on TWAS results from A) all brain tissue panels, and B) each panel separately.

| Gene Set | Estimate | SE<sup>a</sup> | T<sup>b</sup> | P<sup>c</sup> | P.CORR<sup>d</sup> | Panel |
| --- | --- | --- | --- | --- | --- | --- |
| KEGG.LONG.TERM.POTENTIATION | 0.77 | 0.19 | 4.17 | $1.53 \times 10^{-5}$ | $4.98 \times 10^{-2}$ | All brain |
| GO.DENDRITIC.SHAFT | 1.06 | 0.26 | 4.11 | $1.98 \times 10^{-5}$ | $4.98 \times 10^{-2}$ | All brain |
| GO.MEMORY | 0.61 | 0.15 | 4.09 | $2.17 \times 10^{-5}$ | $4.98 \times 10^{-2}$ | All brain |

*B) Panel-specific*
| Gene Set | Estimate | SE<sup>a</sup> | T<sup>b</sup> | P<sup>c</sup> | P.CORR<sup>d</sup> | Panel |
| --- | --- | --- | --- | --- | --- | --- |
| GO.MACROMOLECULAR.COMPLEX.BINDING | 0.46 | 0.09 | 5.09 | $1.83 \times 10^{-7}$ | $5.53 \times 10^{-4}$ | GTeX Caudate basal ganglia |
| GO.MICROTUBULE.BINDING | 0.98 | 0.22 | 4.45 | $4.39 \times 10^{-6}$ | $6.64 \times 10^{-3}$ | GTeX Caudate basal ganglia |
| GO.ALCOHOL.BINDING | 1.75 | 0.38 | 4.58 | $2.32 \times 10^{-6}$ | $7.69 \times 10^{-3}$ | GTeX Pituitary |
| GO.PROTEIN.COMPLEX.BINDING | 0.45 | 0.11 | 4.19 | $1.37 \times 10^{-5}$ | $1.38 \times 10^{-2}$ | GTeX Caudate basal ganglia |
| GO.CHROMATIN.BINDING | 0.77 | 0.19 | 4.12 | $1.91 \times 10^{-5}$ | $1.45 \times 10^{-2}$ | GTeX Caudate basal ganglia |
| GO.LIGAND.DEPENDENT.NUCLEAR.RECEPTOR.BINDING | 1.92 | 0.50 | 3.87 | $5.44 \times 10^{-5}$ | $3.29 \times 10^{-2}$ | GTeX Caudate basal ganglia |

*Note.* Shows results significant after multiple testing correction (P.CORR < 0.05).
<sup>a</sup>Standard error
<sup>b</sup>T-statistic
<sup>c</sup>Raw p-value
<sup>d</sup>Multiple testing-corrected p-value using FDR method

### Comparison with Previous Literature

First, our findings were compared to the largest study of observed gene expression to date (6). Of their significant findings (FDR < 0.1), we replicated five unique genes *(PAPPA2, MBNL1, TMEM64, GNPTAB,* and *KTN1),* pertaining to nine features, with a concordant direction of effect for three genes *(PAPPA2, GNPTAB, KTN1)* (Supplementary Table S7, Supplementary Text).

To evaluate the consistency and novelty of our findings, we compared our results with three previous studies of predicted gene expression changes in MD, performed by Wray et al. (5), Gaspar et al. (12), and Gerring et al. (13). Our results were highly concordant with previous TWASs, as shown by the substantial overlap in the genes found, and high correlations of *z-* scores (*r* ranging from 0.81 – 1) (Supplementary Text, Supplementary Table S8). Moreover, across all TWASs performed to date, we identified 50 novel associations (unique genes). This might be due to differences in the type and number of SNP-weight sets used (e.g. whole blood from the NTR vs. depression genes and networks cohort), methods (e.g. FUSION vs. PrediXcan), and statistical thresholds (e.g. permutation-based vs. Bonferroni statistical threshold).

## Discussion

This is the first study to uncover the *cis-*genetic component of gene expression in major depression through *(i)* a comprehensive investigation in multiple tissues, and *(ii)* an in-depth characterization of the identified associations in follow-up analyses. Here, we highlight a few key findings. First, we detected 91 unique genes associated with depression, 50 of which were novel, undetected by previous TWASs (5,12,12). Fifty-seven of the significant genes were colocalized, 44 were independent, and eight were identified as high-confidence associations based on strong TWAS association, colocalization and causal fine-mapping.

### Key Findings

When testing for an *association between GE and MD,* we detected 91 transcriptome-wide significant genes, differentially expressed across multiple tissues, thus demonstrating the presence of widespread transcriptomic changes in depression. Comparison with previous literature highlighted the novelty of this study, which enabled the identification of 50 novel genes compared to previous TWASs, and 41 genes outside of genome-wide significant loci in the MD GWAS.

When exploring whether significant associations were driven by pleiotropy or linkage using a *colocalization analysis*, we observed that 57 GE - MD relationships derived from the same causal polymorphisms underlying MD SNP associations. This indicated that most of the detected genes constituted pleiotropic effects as opposed to linkage. While these findings suggest that GE mediates the relationship between genetic susceptibility and depression, colocalization does not test for such relationships, nor can it identify causal variants (10).

With a TWAS fine-mapping approach called FOCUS we estimated the probability that each feature within a locus is causally related to MD and identified sets of genes likely to contain the causal feature. In some instances, FOCUS clearly highlighted a single feature as the causal association, such as the upregulation of *NEGR1* in the blood, and downregulation of *PROX2* in the thyroid.

Further investigation of significant associations through a *conditional analysis* determined whether gene-associations within the same genomic region represent independent associations, or whether multiple genes are associated due to correlated predicted expression. The 91 unique significant genes represented 44 independent associations with depression. The strength of association for each feature can be affected by different characteristics of the SNP-weight sets (e.g. sample size), thus warranting cautionary interpretations of which features are driving the associated loci. Comparison of the GWAS summary statistics before and after conditioning on significant TWAS associations additionally revealed that *GWAS associations were explained to a major extent by TWAS associations,* further suggesting the possibility of transcriptomic mediation of genetic risk for depression.

Of the eight high-confidence associations identified, three should be highlighted due to their functional role: the Neuronal Growth Regulator 1 *(NEGR1),* the Estrogen Receptor 2 *(ESR2),* and the Transmembrane Protein 106B *(TMEM106B)*. SNPs within these three genes have been previously detected in one or more GWASs of depression (4,5,27). This study contributed to previous literature by elucidating the functional characteristics of such genes, showing upregulation for *NEGR1* and *TMEM106B,* and downregulation for *ESR2.*

*NEGR1* plays a role in axon extension, synaptic plasticity, and synapse formation, processes key to neuronal functioning (28–30). Moreover, it includes variants found to be related to obesity, a trait repeatedly correlated with MD, both at phenotypic and biological levels (Milaneschi et al., 2019). *ESR2* regulates the activity of estrogen, a sex hormone involved in HPA axis activity and inflammation (31,32). Estrogen fluctuations across women’s lifespan, for example during premenstrual monthly period and menopause, have been proposed as a risk factor for depression (33). However, it remains unclear whether the risk is driven by estrogen or other co-occurring hormonal changes (33,34). *TMEM106B* was previously implicated as a susceptibility gene for neurodegenerative disorders (35–37) and TAR DNA-binding protein 43 (TDP-43) abnormalities (36), which are featured in such pathologies (36,38). Depression has been repeatedly suggested as a risk factor for neurodegenerative disorders (39–41). Moreover, TDP-43 proteinopathy was shown in depression, albeit in a small sample of late-life patients, (42). Overall, while previous literature points to an important role of these genes in depression and related phenotypes, replication of associations is necessary as well as greater insight into the relationship between estrogen levels and TDP-43 with MD.

TWAS-based gene set enrichment analysis was able to identify several gene sets showing dysregulated expression in the brain of individuals with MD. The enriched gene sets are congruent with a previous GWAS-based enrichment analysis (5), corroborating the importance of genes bound by transcription factors RBFOX1/2/3 and FMRP, genes associated with schizophrenia, genes encoding synaptic proteins, and genes encoding ion channels. Three novel gene-sets, here identified as dysregulated across the brain, but not previously found using GWAS-based enrichment analysis, are long term potentiation, dendritic shaft, and memory gene sets. Further investigation of depression-associated differential expression in these gene sets is warranted.

### Limitations of the present study and suggestions for future research

While these findings are promising, several limitations merit discussion. First, the *small sample sizes* of the GE reference samples may have impeded the detection of subtle effects of the transcriptome on depression, meaning that larger samples are needed.

Second, we analysed a wider set of tissues than previous MD TWAS studies, with *20 distinct SNP-weight sets* from blood, brain, HPA and HPT. This may uncover more true associations, but may also have introduced noise since utilizing tissues not strictly relevant to depression might capture non-causal genes (43). An alternative strategy would be use data-driven tissue selection, leveraging an LD score regression-based method (44).

Furthermore, our TWAS approach, by *solely assessing the cis-genetic component of GE,* cannot capture global GE changes nor *trans*-eQTL effects. Studies of observed GE are necessary to examine the former. Future research should additionally channel resources towards building larger GE reference panels to enable investigation of *trans-*eQTL effects.

Lastly, whilst this study provided further insight into *the relationship between SNPs, gene expression, and depression,* and used colocalization and causal fine-mapping analyses to test certain criteria of a causal model, it cannot prove causality between associated genes and depression.

In conclusion, we provided evidence for widespread transcriptomic changes in MD. Our study enabled the detection of novel associations and the elucidation of the transcriptomic changes which previously identified risk genes undergo. We have underlined genes that might be of key relevance to depression, including *NEGR1, ESR2,* and *TMEM106B.* These results suggested an important role of the genetic component of GE in depression.

### URLs

TWAS FUSION SNP-weight sets: http://gusevlab.org/projects/fusion/#reference-functional-data TWAS FUSION website: http://gusevlab.org/projects/fusion/

Frequently asked questions about the TWAS FUSION method: http://twas-hub.org/about/

## Acknowledgments

We thank Alexander Gusev for assisting us with the interpretation of the conditional analysis findings and who created the TWAS FUSION method used here. This paper represents independent research funded by the MRC (MR/N015746/1), and the National Institute for Health Research (NIHR) Biomedical Research Centre at South London and Maudsley NHS Foundation Trust and King’s College London. The authors acknowledge use of the research computing facility at King’s College London, Rosalind (https://rosalind.kcl.ac.uk), which is delivered in partnership with the NIHR Maudsley BRC, and part-funded by capital equipment grants from the Maudsley Charity (award 980) and Guy’s & St. Thomas’ Charity (TR130505). The views expressed are those of the authors and not necessarily those of the NHS, the NIHR or the Department of Health and Social Care. We thank the research participants and employees of 23andMe for making this work possible.

