## Supplementary Text for "Delineating the Genetic Component of Gene Expression in Major Depression"

### Datasets

The Major Depression GWAS summary statistics were obtained from the Psychiatric Genomics Consortium (<https://www.med.unc.edu/pgc/>) and converted to an LD-score format for readability by the FUSION software, employed in the TWAS analyses. The adaptation was performed with the LDSC munge_stats.py utility using Python (version 3.5.1).

### Statistical analyses

#### Calculation of the transcriptome-wide significance threshold

Firstly, as expected in the FUSION protocol, GE levels were inferred for all heritable features (*N* = 83,002) based on the selected SNP-weight sets and the 1000 Genomes LD reference data. GWAS summary statistics were not used as the calculation of the significance threshold should be based on a null phenotype. A thousand permutations were performed, with each permutation randomly generating a normally-distributed null phenotype. Such phenotype was subsequently tested in association with predicted GE levels. Given the use of a random phenotype, the identified feature – trait relationships should constitute false-positive findings, uniquely attributable to chance. To develop a significance threshold able to distinguish between chance and meaningful findings, the minimum *p-*value of all individual permutations (*N_permutations_* = 1,000) was collected to form a normal distribution of minimum *p-*values. The five percent quantile of this distribution, equalling to a false positive rate of *α* = .05 (1), was considered as our transcriptome-wide significance threshold, which corresponded to *p* < 2.13×10^-6^. We additionally calculated a more stringent threshold to capture genes of high significance (*α* = 0.001): *p* < 4.38×10^-8^.

#### TWAS FUSION

Gene expression-depression associations were obtained only for features with a non-zero *cis-*SNP heritability (*p* < 0.01). This was calculated with the Average Information Restricted Maximum Likelihood (AI-REML) algorithm of Genome-wide Complex Trait Analyses (GCTA). SNPs were selected if they resided within the gene boundaries ±500kb.

A TWAS FUSION analysis was run for all significantly heritable features, for each feature, based on one of several predictive models (BLUP, LASSO, elastic net, or BSLMM), with the best-fitting model being used. TWAS *Z*-scores were estimated, for each feature separately, based on the following linear model:

Feature *Z*-score = *w_1_z_1_* + *w_2_z_2_* + *w_3_z_3_* + *w_4_z_4_* …

Where *w_i_* = the correlation of a SNP within a feature with the gene expression of such feature,

*z_i_* = the standardized effect from the GWAS which tested the association between that SNP and a trait, and

*i* = the SNP within the feature.

Outputs from all chromosomes and SNP-weight sets were merged and subsequently filtered based on our transcriptome-wide significant threshold.

#### Colocalization

This method used a Bayesian approach estimating the posterior probability (PP) of five models concerning GWAS and TWAS associations. These involve a SNP being in:

PP0) No association with depression or GE (null findings),

PP1) Association with depression only (GWAS significance),

PP2) Association with GE only (TWAS significance),

PP3) Association with both, from two independent SNPs (GWAS and TWAS significance, two SNPs involved),

PP4) Association with both, at a shared SNP (GWAS and TWAS significance, one SNP only).

By comparing values for models three and four (PP3 vs. PP4), we can distinguish whether the GWAS and TWAS associations are colocalized, i.e. whether the signal for an association with the trait and with GE at a locus results from the same causal polymorphism. To perform colocalization, the *coloc* R package (2), available in the FUSION software, was employed. Colocalization was implemented only for genes surpassing transcriptome-wide significance.

#### Conditional Analysis

With a conditional analysis, we could identify loci of co-expression as well as distinguish between independent and conditioned features. *Independent/jointly significant features* remain associated with the phenotype, at a nominal significance level (*p* < 0.05), after adjustments. *Conditioned* or *marginally significant features* are those whose association with depression is solely reliant on the expression of other nearby features. Conditioned features are thus significantly associated with depression in the *un*adjusted model only.

A conditional analysis can additionally show to what extent GWAS signal is attributable to functional associations. In fact, in such analysis, the top SNP in a locus is conditioned on the GE patterns of the most significant feature in the same locus. Subsequently, the variance in a SNP signal accounted for by such functional associations was calculated, with the following formula:

*R^2^* = 1 – *χ^2^ _conditioned GWAS association_* / *χ^2^ _unconditioned GWAS association_*

Where *R^2^* = variance explained,

*χ^2^ _conditioned GWAS association_* = SNP-phenotype association after adjusting for the GE of the most significant feature in the locus,

*χ^2^ _unconditioned GWAS association_* = SNP-phenotype association before adjusting.

The conditional analysis was performed for chromosomal regions with multiple statistically significant features, within and across SNP-weight sets, as described in the FUSION webpage (<http://gusevlab.org/projects/fusion/>). Features were defined as pertaining to a shared locus when boundaries overlapped within ±1.5Mb.

#### TWAS-GSEA

For the TWAS-GSEA, a linear mixed model was run. Gene set membership was regressed on the TWAS feature *z*-score indicating non-zero association. Gene set membership was developed as a dichotomous variable with information on whether a gene pertains to a given functional annotation. Analyses were performed with the R package lme4qtl (3), which permits the fitting of linear mixed models.

We firstly conducted a *hypothesis-free* TWAS-GSEA testing all annotations from the gene ontology (GO) resource, a database where gene and gene set functions are specified. *Candidate gene sets* were additionally tested. These contained pathways previously selected by Wray et al. (4) due to their involvement in psychiatric disorders. We attempted to validate these with a functionally-informed gene set enrichment analysis.

### Results

#### Comparison with previous literature

##### Literature of observed gene expression

We compared our results with the Jansen et al (5) publication, which assessed whole blood GE using microarray technology in the Netherlands Study of Depression and Anxiety. We replicated five unique genes (*PAPPA2, MBNL1, TMEM64, GNPTAB,* and *KTN1*), from nine features. Of these, *PAPPA2* (GTEx Substantia nigra) was upregulated in both studies. Contrarily, *KTN1* (CMC DLPFC Splicing, GTEx Cerebellum), *MBNL1* (CMC DLPFC), and *GNPTAB* (GTEx Thyroid) were consistently downregulated. A contrasting direction of effect between the two studies was present for *TMEM64*, *MBNL1* (CMC DLPFC Splicing only), and *GNPTAB* (GTEx Caudate).

##### Literature of predicted gene expression

We compared our findings with three previous TWAS of major depression. Overall, a total of 133 unique genes, significant in *any* SNP-weight set, in *either* of the four TWASs, were considered.

As for the Wray et al. (4) findings, within the CMC DLPFC, i.e. the only SNP-weight set they tested, we observed identical *z*-scores (*r_z-scores_ =* 1), as expected. Yet, due to our greater multiple testing burden, we did not identify as many genes as significant within the CMC DLPFC (we tested for 20, as opposed to one, SNP-weight sets). Contrarily, because of the greater variety of weights being tested, overall, we uncovered more associations (177 vs. 17).

When comparing our findings to Gaspar et al. (6), similar results were found (*r_z-scores_* = 0.81). Of the 25 unique genes significantly associated with MD in the Gaspar et al. (6) study, we failed to detect 12 as significant. Of our 91 significant unique genes, Gaspar et al. (6) did not identify 78 associations.

Finally, when contrasting our results to Gerring et al.’s (7), we also observed similar z-scores (*r_z-scores_* = 0.85). In their TWAS, Gerring et al. (7) identified 57 unique genes in association with depression, at a Bonferroni significance level. Of such genes, we failed to detect 29 associations. On the other hand, we presented 63 significant genes that Gerring et al. (7) did not previously uncover. Overall, across all TWASs, we unraveled 50 novel associations.

Supplementary Figures

a.
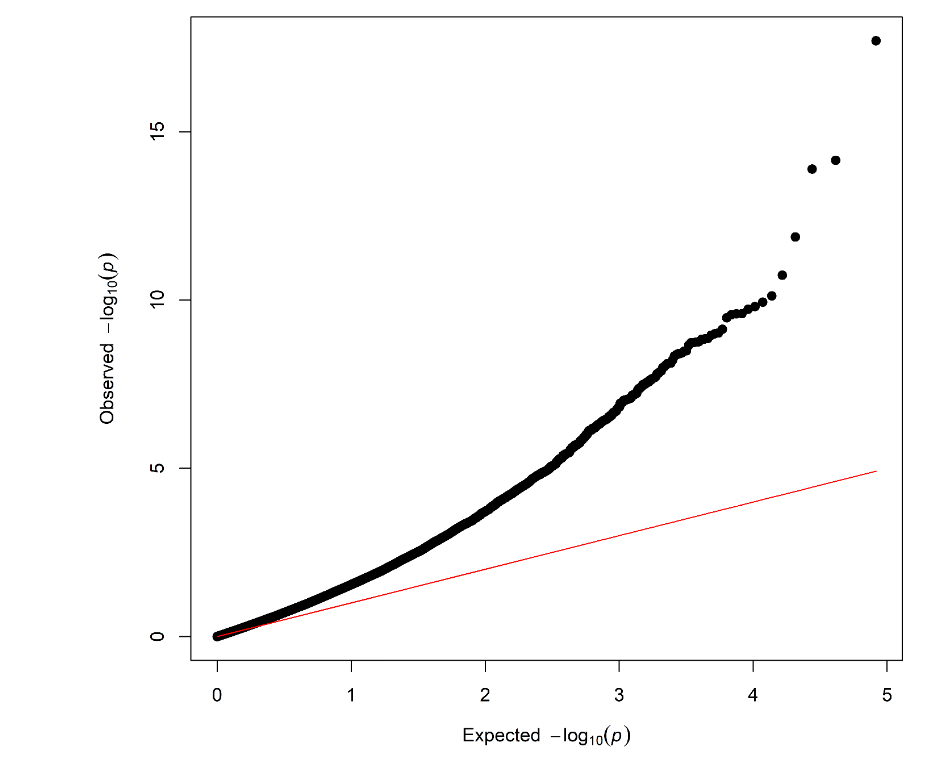
b.
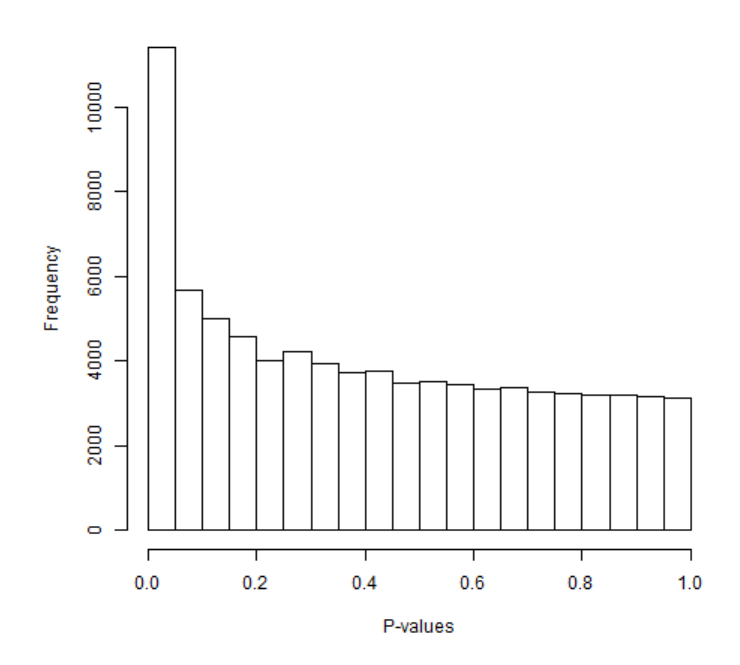

*Supplementary Figure S1a-b.* The relationship between major depression and the genetic component of gene expression. (a) QQ-plot showing the distribution of *p-*values expected under the null hypothesis (red line) versus the observed distribution (black line). Inflation is observed. In line with the Wray et al. (2018) GWAS results, this is likely the result of polygenicity as opposed to linkage disequilibrium. (b) Histogram of *p-*values: the equal distribution of *p-*values at the bottom represents the null hypothesis being met. The peak in correspondence to the smallest *p-*values provides evidence for our alternative hypothesis of an association between depression and GE.
